# *Pseudomonas arenae* sp. nov., *Pseudomonas glycinis* sp. nov. and *Pseudomonas harudinis* sp. nov., three novel bacterial species and plant endophytes

**DOI:** 10.1101/2021.05.13.444027

**Authors:** Sarah Seaton, Jacqueline Lemaire, Patrik Inderbitzin, Victoria Knight-Connoni, James F. White, Martha E. Trujillo

## Abstract

Three novel *Pseudomonas* species associated with healthy plants are described from the United States. They are *Pseudomonas arenae* sp. nov. from soybean in Missouri and *Phragmites* sp. in New Jersey; *Pseudomonas glycinis* sp. nov. from *Vaccinium macrocarpon* fruit in Massachusetts, groundwater in Tennessee and soybean in Indiana; and *Pseudomonas harudinis* sp. nov. from *Phragmites* sp. in New Jersey. No pathogenic strains are known for any of the novel species based on genome comparisons to assemblies in GenBank.

## INTRODUCTION

*Pseudomonas* is a large and diverse genus currently comprising more than 250 named species, which are phenotypically and genotypically well-defined (Parte et al. 2020). These organisms belong to the gamma subclass of Proteobacteria, are rod-shaped, Gram-type negative chemoheterotrophs that are motile by means of polar flagella. *Pseudomonas* species have simple nutritional requirements and can utilize an array of small organic molecules as sources of carbon and energy. Such nutritional diversity is reflected in the relative abundance of these organisms in nature, with as many as 10^6^ fluorescent pseudomonads residing in a single gram of soil (Tarnawski et al. 2003; Vančura 1980). *Pseudomonas* species are strictly aerobic, however, some can utilize NO_3_ as an electron acceptor in place of O_2_. In addition, nitrate can be used as a nitrogen source for all known species (Stanier et al. 1966).

Pseudomonads are ubiquitous in the rhizosphere, often living in a commensal relationship with plants, utilizing plant-exuded nutrients and occupying sites provided by the architecture of the plant. Such commensal species, in turn, impact plant health by suppressing phytopathogens (Couillerot et al. 2009; Gerbore et al. 2014), enhancing local access to nutrients, and inducing systemic resistance in the plant host (Bakker et al. 2007; De Vleesschauwer et al. 2008). A subset of rhizosphere pseudomonads are further adapted to live as endophytes, residing within cells, the intercellular spaces or the vascular system of host plants (Mitter et al. 2013). Here, we describe three such endophytic strains, representing three novel species of the genus *Pseudomonas. Pseudomonas arenae* sp. nov. strain VK110 and *Pseudomonas harudinis* sp. nov. strain SS112 were isolated from *Phragmites* reed grass, collected roadside in New Jersey, USA. *Pseudomonas glycinis* sp. nov. strain PI110 originated from healthy field-grown soybean in Indiana, USA. We provide phenotypic and phylogenomic details describing these three species.

## METHODS

### Isolation

Strain PI111 was isolated from healthy field-grown *Glycine max* in Indiana, United States, and strains SS112 and VK110 from *Phragmites* sp. in New Jersey, United States. Plant tissue was washed with a mild detergent to remove particulates, surface-sterilized with bleach (1% v/v sodium hypochlorite) and ethanol (70% v/v), and homogenized. Serial dilutions of tissue homogenate were plated on a panel of media types for endophyte cultivation. All strains were streaked to purity and stored in glycerol (20% v/v) at −80°C until subjected to further testing.

### Motility

The strains were tested for flagellar-dependent swimming and swarming motility on R2A plates solidified with 0.3% and 0.6% agar, respectively. Three independent colonies were inoculated onto R2A broth and grown for 36 hr at 24°C. Broth cultures were normalized to an OD600 of 0.1, and 1.5 µl of culture was spotted directly onto the surface of the motility agar. The diameter of colony expansion was measured for 5 days.

### Carbon source utilization

Substrate utilization was assessed using Biolog GenIII Microplates (Catalogue No. 1030) (Biolog Inc., Hayward, CA). Each bacterium was inoculated in duplicate plates using Protocol A, described by the manufacturer, with the exception that plates were incubated at 30°C. Respiration leading to reduction of the tetrazolium indicator was measured by absorbance at 590 nm.

### Biochemical analyses

Catalase activity was evaluated by immediate effervescence after the application of 3 % (v/v) hydrogen peroxide solution via the tube method, a positive reaction was indicated by the production of bubbles. *Staphylococcus aureus* NCIMB 12702 and *Streptococcus pyogenes* ATCC 19615 were used as positive and negative controls, respectively. Oxidase activity was evaluated via the oxidation of Kovács oxidase reagent, 1% (w/v) tetra-methyl-p-phenylenediamine dihydrochloride in water, via the filter-paper spot method. A positive reaction was indicated when the microorganisms color changed to dark purple. *Pseudomonas aeruginosa* NCIMB 12469 and *Escherichia coli* ATCC 25922 were used as positive and negative controls, respectively. Urease activity was evaluated via the hydrolysis of urea in Christensen’s Urea Agar, using phenol red as a pH indicator. *Proteus hauseri* ATCC 13315 and *Escherichia coli* ATCC 25922 were used as positive and negative controls, respectively. Gram staining was performed using standard protocols.

### Phylogenetic and genomic analyses

DNA was extracted from pure cultures using the Omega Mag-Bind Universal Pathogen Kit according to manufacturer’s protocol with a final elution volume of 60µl (Omega Biotek Inc., Norcross, GA). DNA samples were quantified using Qubit fluorometer (ThermoFisher Scientific, Waltham, MA) and normalized to 100 ng. DNA was prepped using Nextera DNA Flex Library Prep kit according to manufacturer’s instructions (Illumina Inc., San Diego, CA). DNA libraries were quantified via qPCR using KAPA Library Quantification kit (Roche Sequencing and Life Science, Wilmington, MA) and combined in equimolar concentrations into one 24-sample pool. Libraries were sequenced on a MiSeq using pair-end reads (2×200bp). Reads were trimmed of adapters and low-quality bases using Cutadapt (version 1.9.1) and assembled into contigs using MEGAHIT (version 1.1.2) (Mitter et al. 2013). Reads were mapped to contigs using Bowtie2 (version 2.3.4) (Langmead and Salzberg 2012), and contigs were assembled into scaffolds using BESST (2.2.8) (Sahlin et al. 2014). Phylogenomic trees were generated using GToTree (version 1.2.1) (Lee 2019).

16S rRNA gene sequences were extracted from genome assemblies using barrnap (Seemann 2019), and 16S rRNA gene phylogenetic analyses were performed using FastTree (Price et al. 2010) using a General Time Reversible substitution model. Taxon sampling for each species is described in the respective phylogenetic tree figure legend.

Average nucleotide identity (ANI) analyses between genome assemblies were performed using the pyani ANIm algorithm (Richter and Rosselló-Móra 2009) implemented in the MUMmer package (Kurtz et al. 2004) retrieved from https://github.com/widdowquinn/pyani.

Geographic distribution and host range of novel species were inferred by ANI to assemblies from unidentified species from GenBank (Ciufo et al. 2018) and the Indigo internal collection. An ANI threshold of ≥95% indicated conspecificity (Chun et al. 2018; Richter and Rosselló-Móra 2009).

## RESULTS

### Phylogenetic and genomic analyses

#### *Pseudomonas arenae* sp. nov. strain VK110

The strain VK110 16S rRNA gene sequence MZ099645 shared 99.7% identity with the 16S of *Pseudomonas rhodesiae* CIP 104664^T^. A phylogenomic tree using GToTree (Lee 2019) confirmed the affiliation of strain VK110 to the genus *Pseudomonas*. VK110 was most closely related to *Pseudomonas rhodesiae* DSM 14020 ^T^ with 100% support (Figure 1). Average nucleotide identity (ANI) to *P. rhodesiae* was 91.2%, well below the threshold for species demarcation (Chun et al. 2018; Richter and Rosselló-Móra 2009), and showing that strain VK110 represents a new genomic species of *Pseudomonas*.

**Figure 1.**
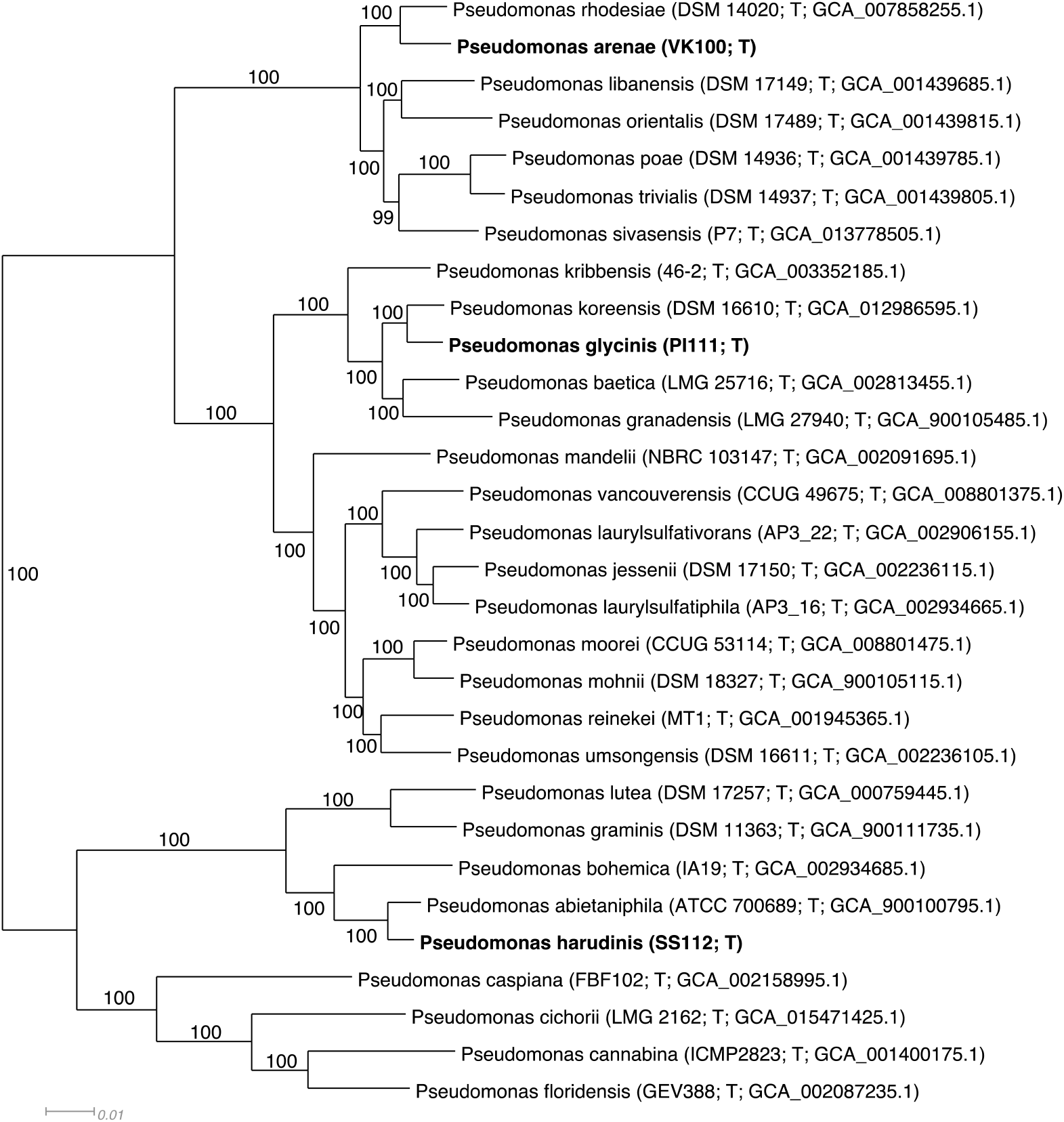
Phylogenomic tree of *Pseudomonas arenae* sp. nov., *Pseudomonas glycinis* sp. nov., *Pseudomonas harudinis* sp. nov. and relatives inferred by GToTree (Lee 2019). Taxon sampling includes a representative set of relatives based on *Pseudomonas* species phylogenies. Alignment consists of 38,520 amino acid positions from 159 – 170 genes depending on taxon. Species described in this study are in bold. Strain numbers and GenBank accession numbers follow species names, T stands for ‘type’. Support values above 50% are given by the branches. *Pseudomonas arenae* is most closely related to *P. rhodesiae, Pseudomonas glycinis* to *P. koreensis* and *Pseudomonas harudinis* to *P. abietaniphila*, all with 100% support. Branch lengths are proportional to the changes along the branches, a scale bar is provided at the bottom.

#### *Pseudomonas glycinis* sp. nov. strain PI111

The strain PI111 16S rRNA gene sequence MZ099646 shared 99.9% identity with the 16S of *Pseudomonas koreensis* KACC 10848 ^T^. A phylogenomic tree using GToTree (Lee 2019) confirmed the affiliation of strain PI111 to the genus *Pseudomonas*. PI111 was most closely related to *Pseudomonas koreensis* DSM 16610^T^ with 100% support (Figure 1). Average nucleotide identity (ANI) to *P. koreensis* was 92.0%, well below the threshold for species demarcation (Chun et al. 2018; Richter and Rosselló-Móra 2009), and showing that strain PI111 represents a new genomic species of *Pseudomonas*.

#### *Pseudomonas harudinis* sp. nov. strain SS112

The strain SS112 16S rRNA gene sequence MZ099647 shared 99.2% identity with the 16S of *Pseudomonas abietaniphila* ATCC 700689^T^. A phylogenomic tree using GToTree (Lee 2019) confirmed the affiliation of strain SS112 to the genus *Pseudomonas*. SS112 was most closely related to *Pseudomonas abietaniphila* ATCC 700689^T^ with 100% support (Figure 1). Average nucleotide identity (ANI) to *P. abietaniphila* was 92.4%, well below the threshold for species demarcation (Chun et al. 2018; Richter and Rosselló-Móra 2009), and showing that strain SS112 represents a new genomic species of *Pseudomonas*.

### Geographic distribution and host range

Geographic distribution and host range of the novel species was inferred by comparison to congeneric genome assemblies of unidentified species from GenBank and the Indigo internal collection. Hits to novel species included the Indigo strain *Pseudomonas arenae* strain JL113 (ANI: 99.2%; query coverage: 95.8%) collected from healthy *Glycine max* plants in Missouri. Hits among genome assemblies from GenBank to *P. glycinis* included assemblies GCF_016756945.1, GCF_017947285.1 and GCF_017947305.1 (ANI: 95.6% - 96.6%; query coverage: 89.9% - 91.7%) from strains originating from the surface of *Vaccinium macrocarpon* berries in Massachusetts, United States, and assemblies GCF_017351195.1, GCF_002901605.1 and GCF_002901465.1 (ANI: 96.9%; query coverage: 94.4 – 94.6%) from strains isolated from groundwater in Tennessee, United States. Known geographic distributions of the novel species from culturing is illustrated in Figure 2, Figure 3 and Figure 4, and substrates are compiled in Table 1.

**Figure 2.**
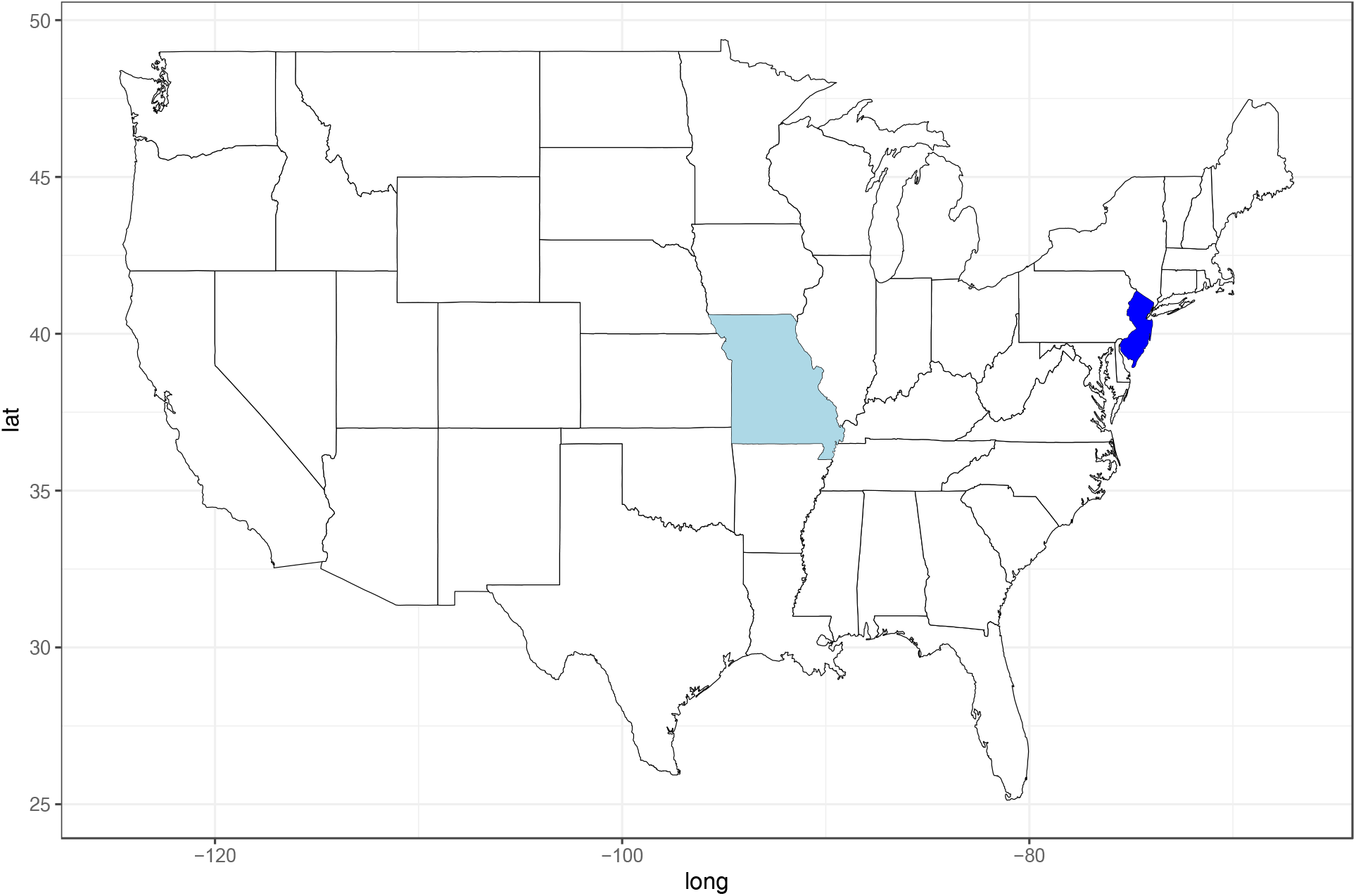
Geographic distribution of *Pseudomonas arenae* sp. nov. based on culturing. Dark blue indicates type strain, light blue additional strains.

**Figure 3.**
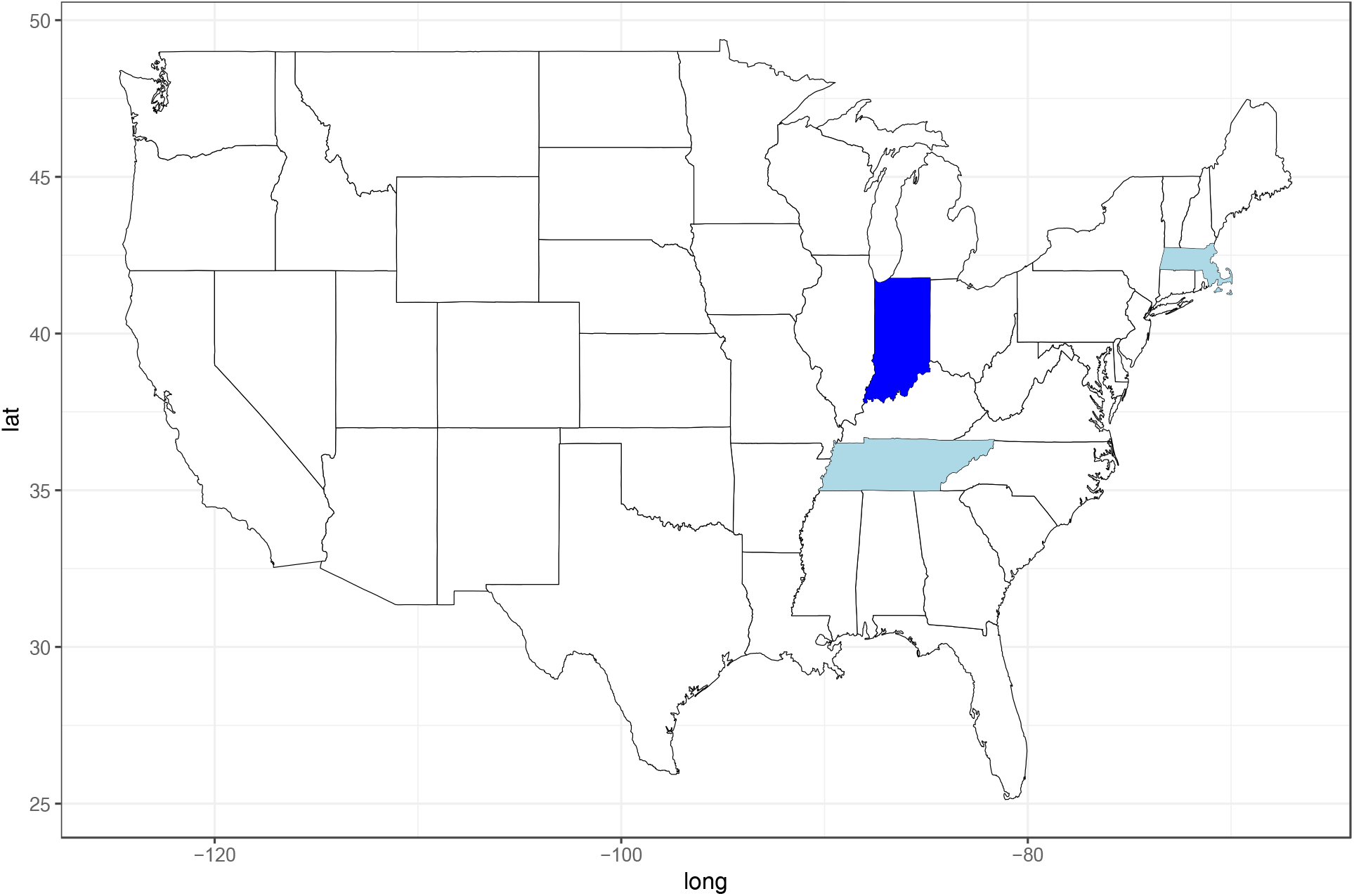
Geographic distribution of *Pseudomonas glycinis* sp. nov. based on culturing. Colors mark states where species was collected. Dark blue indicates type strain, light blue additional strains.

**Figure 4.**
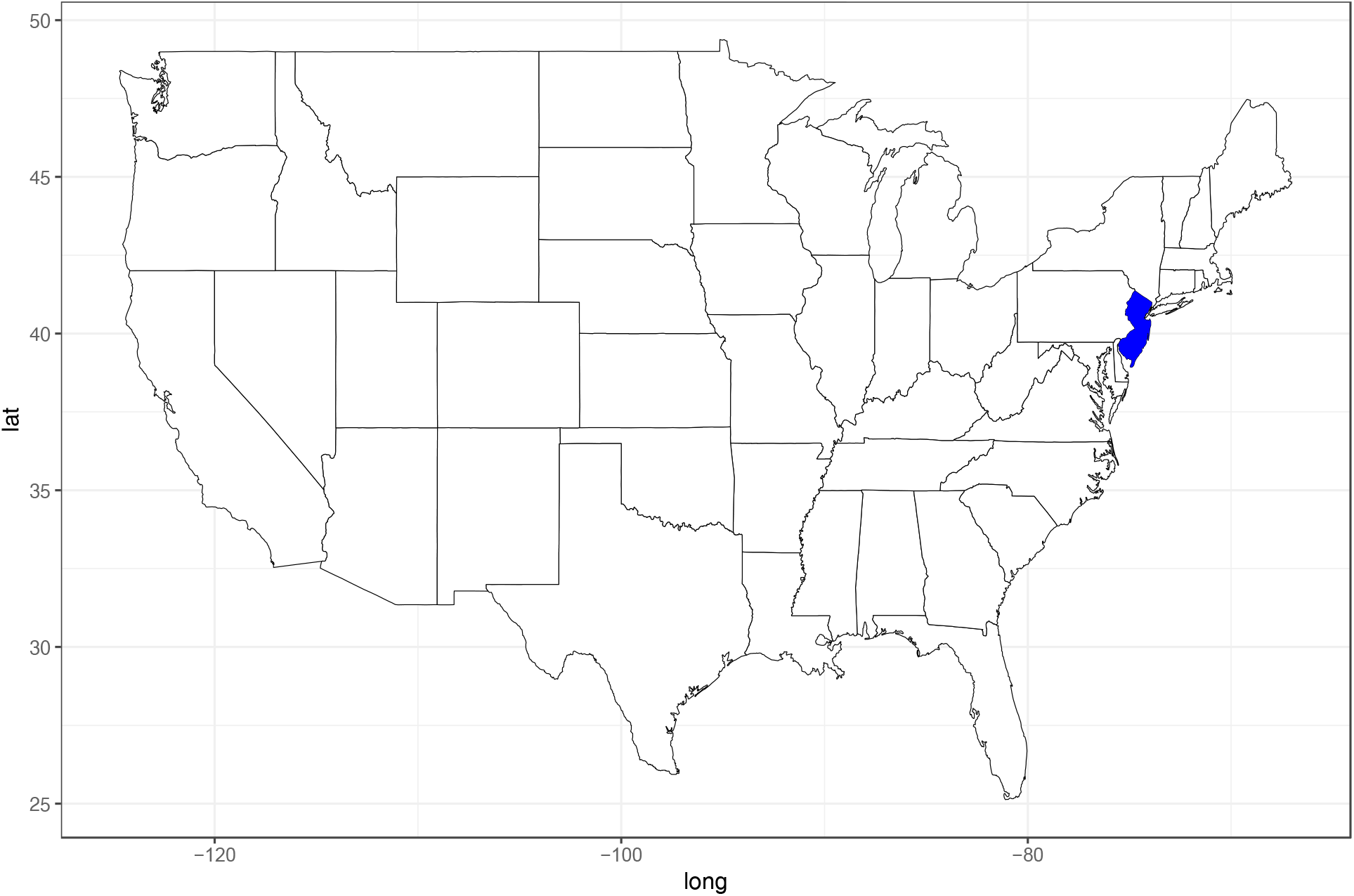
Geographic distribution of *Pseudomonas harudinis* sp. nov. based on culturing. Dark blue marks state where type strain was collected, no cultures are known from other states.

**Table 1.** Substrates of novel species based on culturing.

| **Species** | **Substrate<sup>1</sup>** |
| --- | --- |
| *Pseudomonas arenae* | *Glycine max*, *Phragmites* sp. |
| *Pseudomonas glycinis* | *Glycine max*, *Vaccinium macrocarpon*;<br>groundwater |
| *Pseudomonas harudinis* | *Phragmites* sp. |

<sup>1</sup> No disease symptoms reported for strains isolated from plants.

### Morphology, physiology and biochemical characteristics

The three new isolates studied were Gram-stain negative, non-spore forming, motile rod-shaped organisms that occurred as single cells (Figure 5, Figure 6, Figure 7). All strains also grew well after 24 h on Tryptic soy, R2 and Nutrient agars at 22 and 30°C; strains were inhibited for growth at 37°C. All strains produced circular and smooth colonies. Those of strain PI111 were white and mucoid; light pink colonies were recorded for isolate VK110, and non-pigmented colonies were observed for strain SS112.

**Figure 5.**
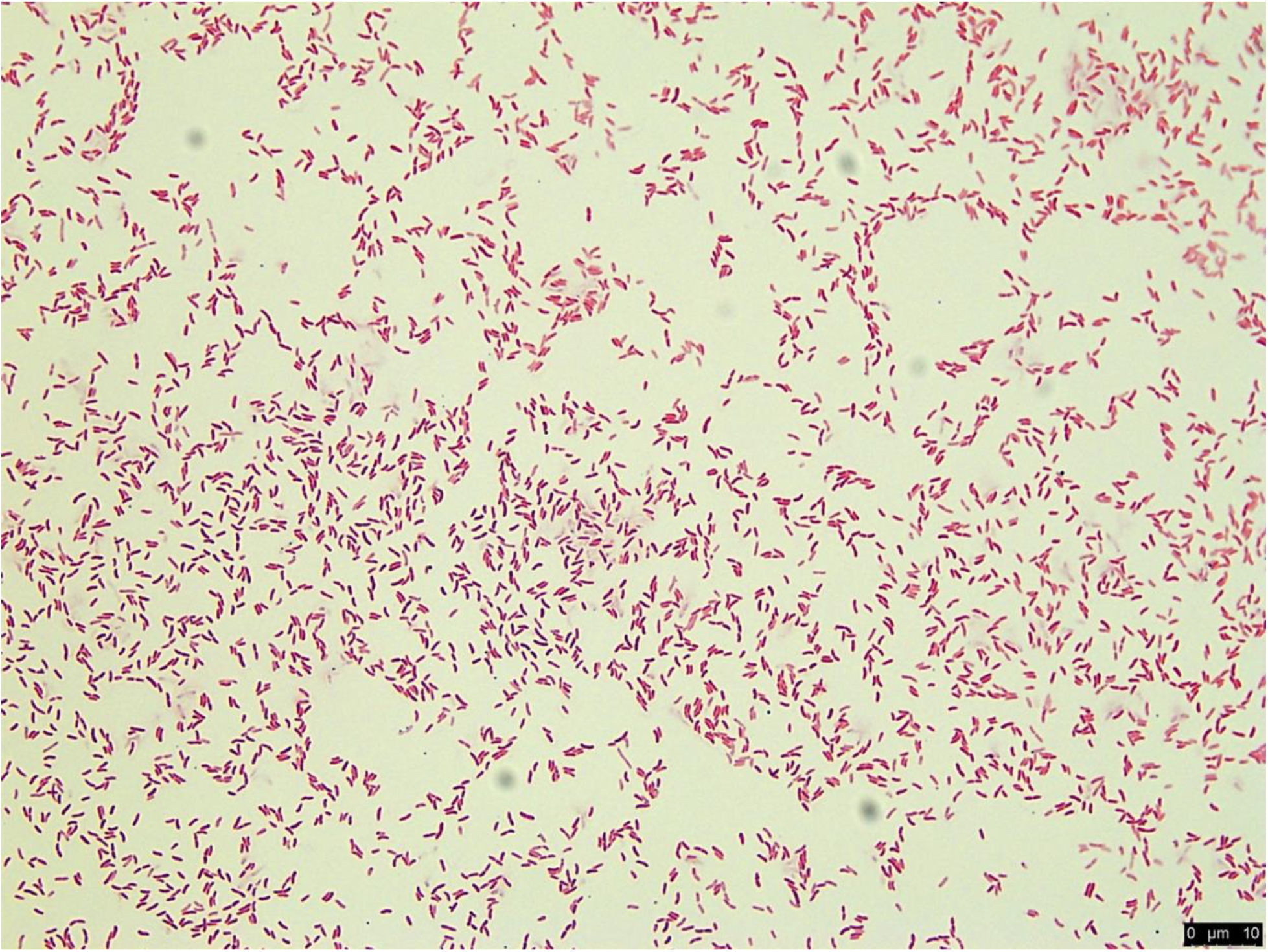
Morphology of *Pseudomonas arenae* sp. nov. strain VK110 depicted following Gram stain under bright field microscopy. Bar = 10 µm.

**Figure 6.**
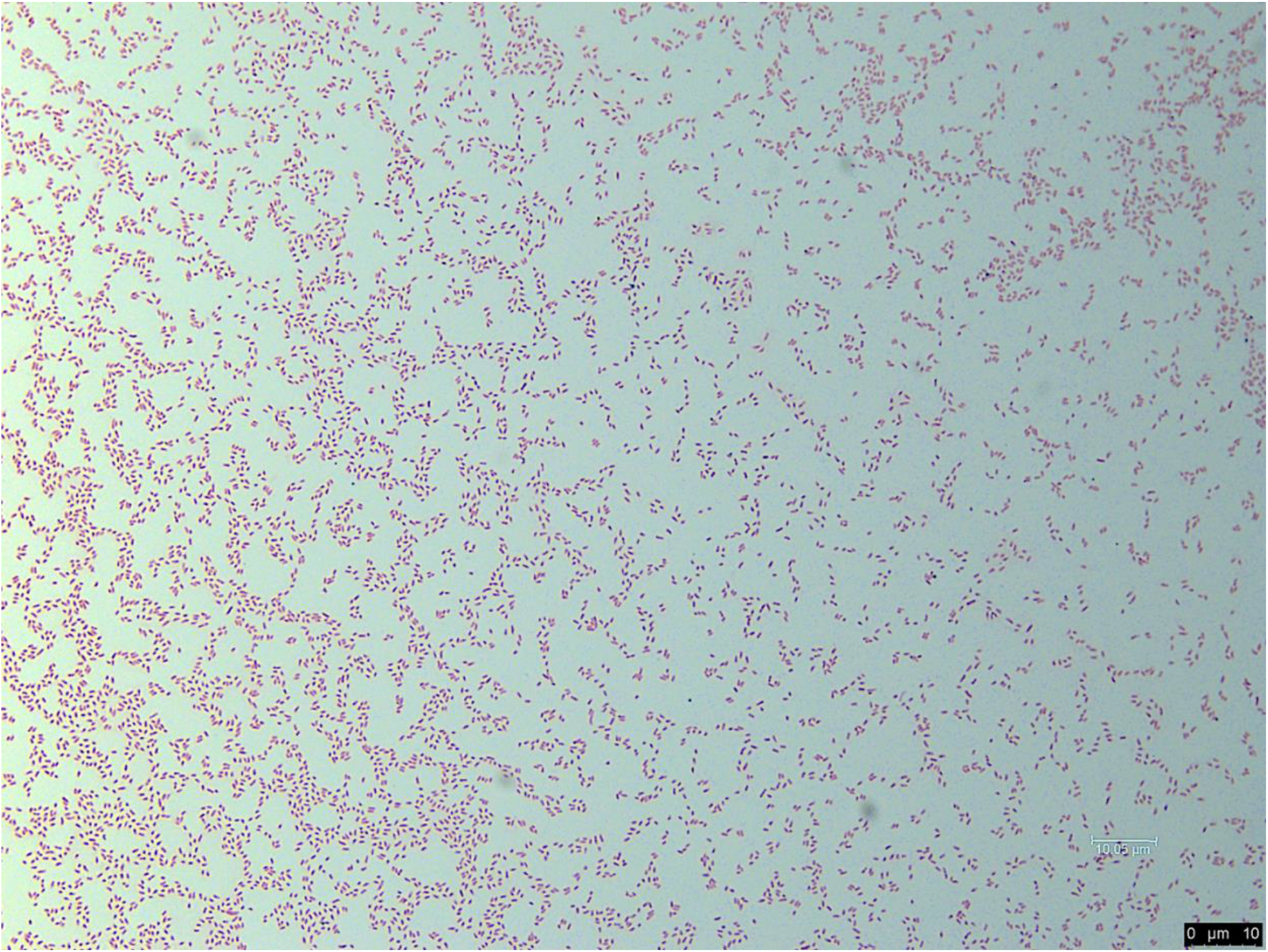
Morphology of *Pseudomonas glycinis* sp. nov. strain PI111 depicted following Gram stain using bright field microscopy. Bar = 10 µm.

**Figure 7.**
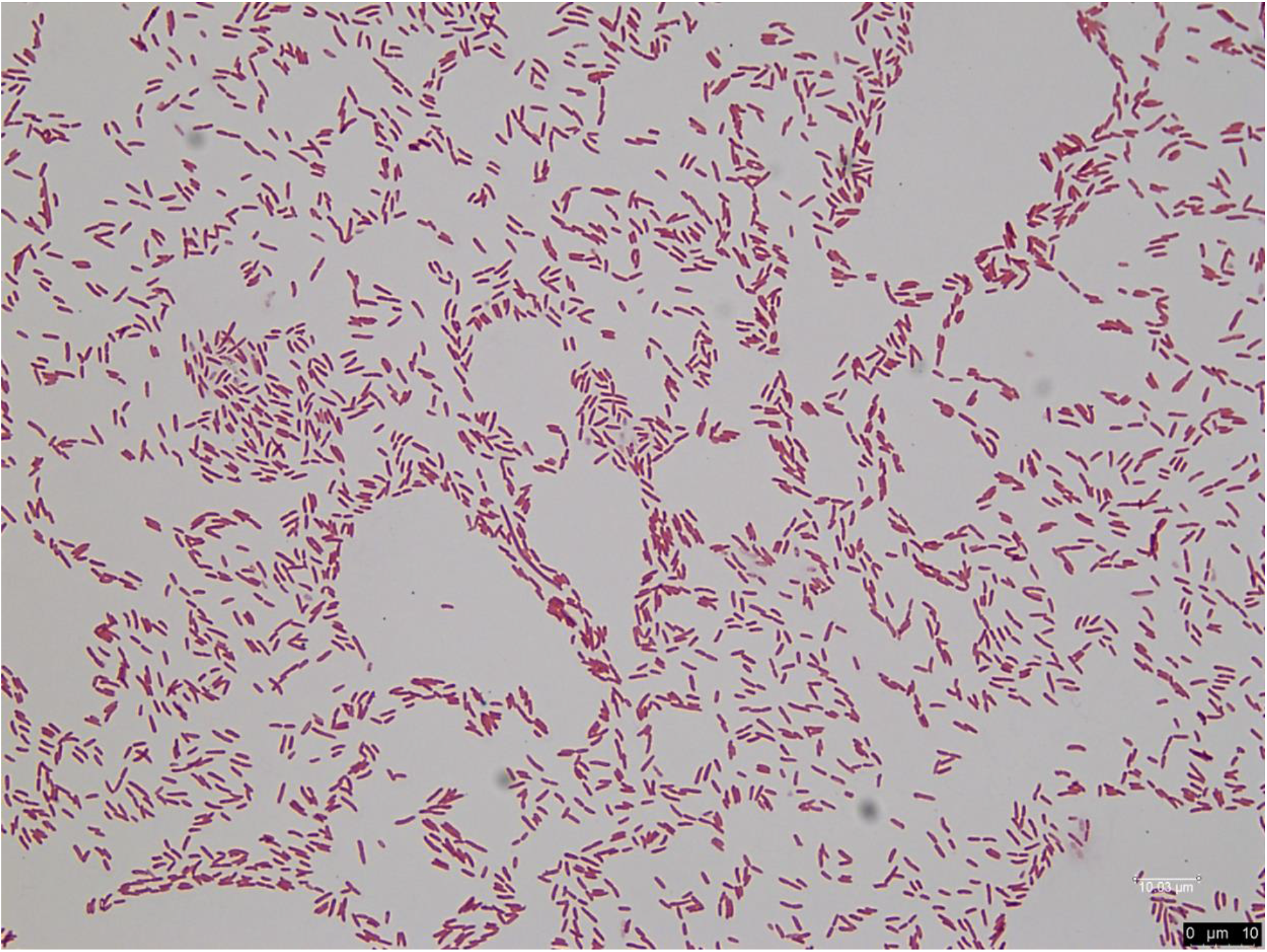
Morphology of *Pseudomonas harudinis* sp. nov. strain SS112 depicted following Gram stain using bright field microscopy. Bar = 10 µm.

All strains were aerobic and positive for catalase activity. In addition, all grew on media supplemented with 1 and 4% NaCl and at pH 5 and 6.

The following substrates were used as carbon sources by all strains when tested on Gen III Biolog Microplates according to the manufacturer’s recommendations: α-D-glucose, D-mannose, D-fructose (weak reaction for SS112), D-galactose, DL-fucose (weak reaction, VK110), inosine (weak reaction, PI111), sodium lactate (1%), fusidic acid (weak reaction for SS112), D-serine (weak reaction for SS112), D-mannitol, D-arabitol (weak reaction, PI111), glycerol (weak reaction, PI111), D-fructose-6-phosphate (weak reaction, VK110 and PI111), L-alanine (weak reaction, PI111), L-arginine, L-aspartic acid (weak reaction, PI111), L-glutamic acid, L-galactonic acid lactone (weak reaction, PI111), D-gluconic acid, glucuronate, mucic acid, quinic acid, D-saccharic acid, L-lactic acid, citric acid, α-keto-glutaric acid, L-malic acid and acetic acid.

None of the strains used the following carbon sources: dextrin, D-maltose, D-cellobiose, gentiobiose, D-turanose, stachyose, D-raffinose, α-D-lactose, D-melibiose, β-methyl-D-glucoside, D-salicin, N-Acetyl-β-D-mannosamine, N-Acetyl-β-D-galactosamine, N-Acetyl neuraminic acid, 2-methyl glucose, D-glucose-phosphate, p-hydroxy-phenylacetic acid, α-hydroxy-butyric acid, α-keto-butyric acid, acetoacetic acid, formic acid and sodium butyrate.

All strains were resistant to the following antibiotics: lincomycin, troleandomycin, rifamycin and aztreonam and grew in the presence of guanidine HCl, tetrazolium violet, tetrazolium blue and potassium tellurite. None of the strains hydrolyze gelatin or pectin.

A set of phenotypic characteristics that differentiate between each one of the new species and their corresponding phylogenetic neighbors is found in Tables 2, 3 and 4, for *P. arenae* sp. nov., *P. glycinis* sp. nov. and *P. harudinis* sp. nov., respectively.

**Table 2.**
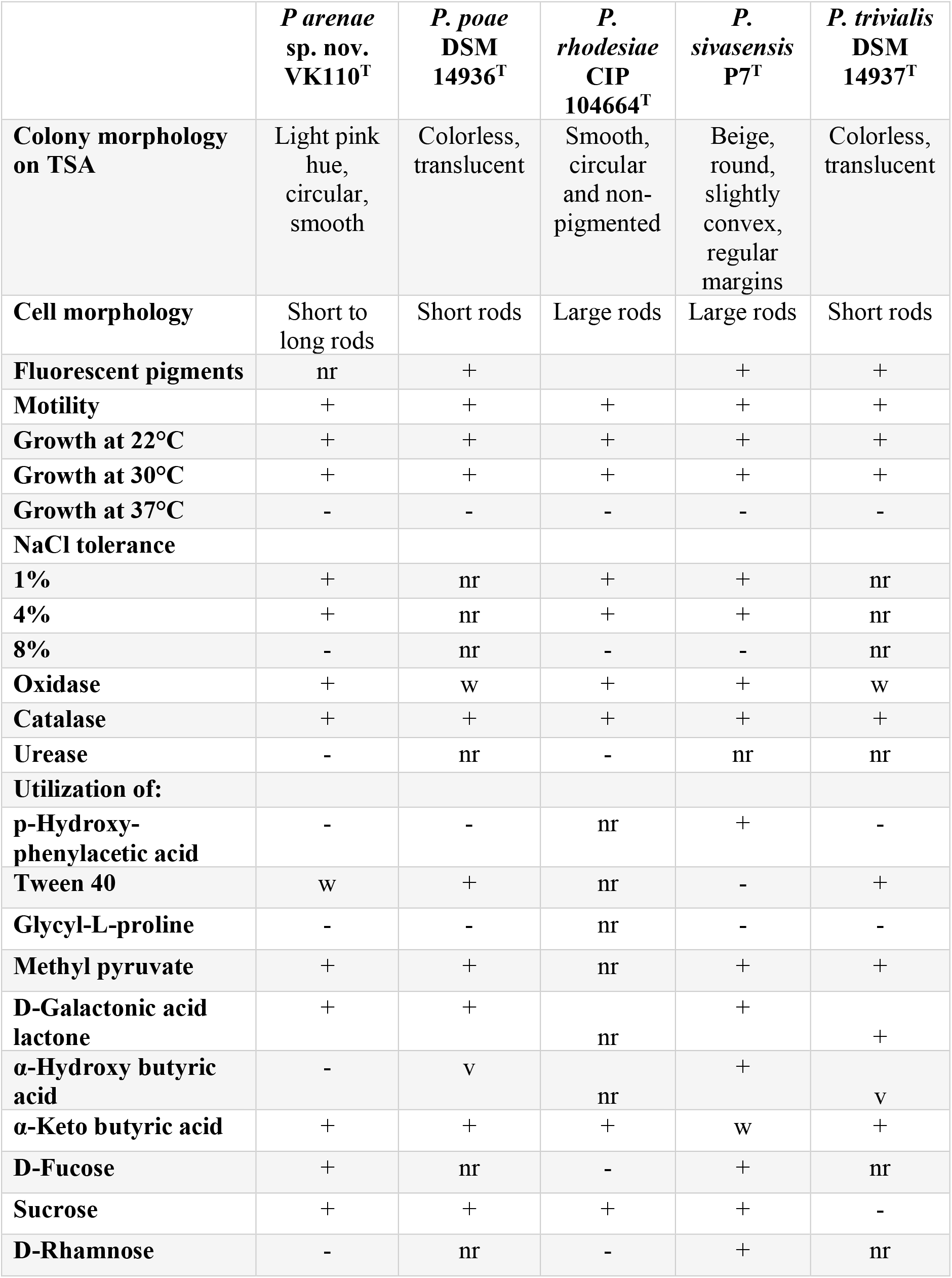
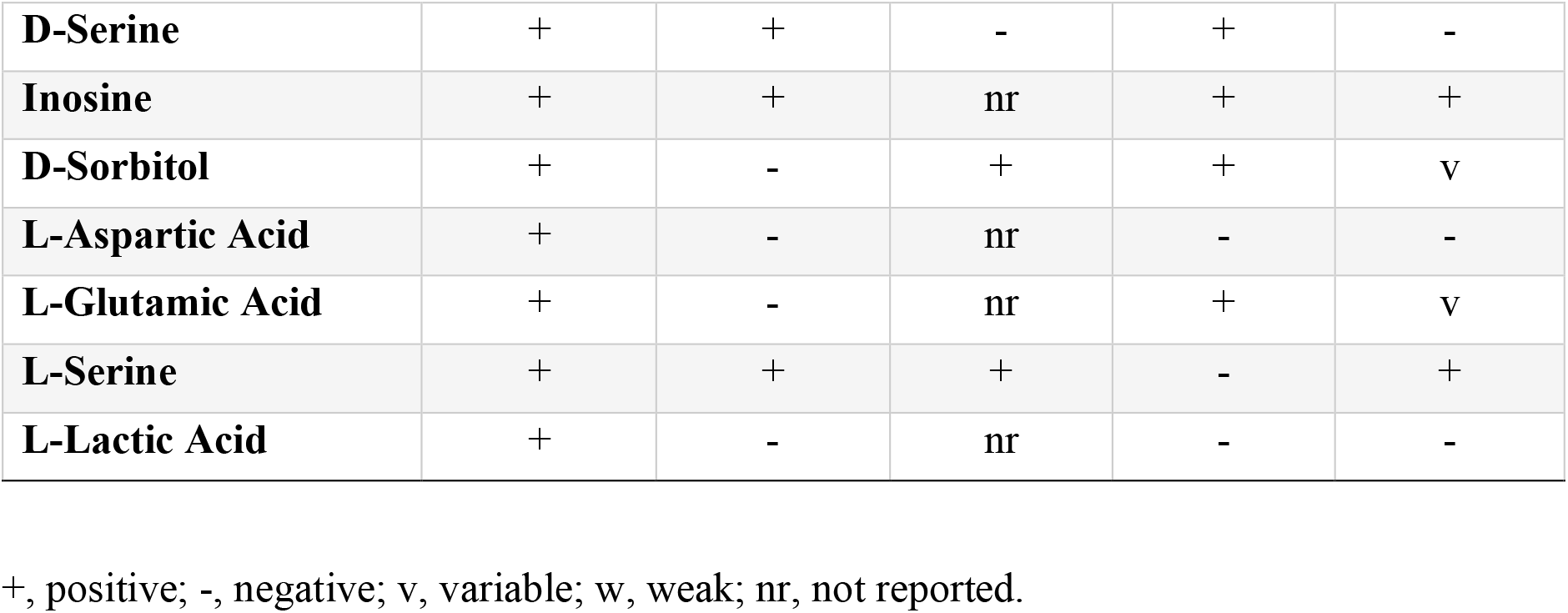
Comparative physiological characteristics among *Pseudomonas arenae* sp. nov. VK110 and related type strains. Data from this study and Coroler et al. (1996); Behrendt et al. (2003); Duman et al. (2020).

|  | *P. arenae*<br>sp. nov.<br>VK110<sup>T</sup> | *P. poae*<br>DSM<br>14936<sup>T</sup> | *P. rhodesiae*<br>CIP<br>104664<sup>T</sup> | *P. sivasensis*<br>P7<sup>T</sup> | *P. trivialis*<br>DSM<br>14937<sup>T</sup> |
| --- | --- | --- | --- | --- | --- |
| **Colony morphology on TSA** | Light pink hue, circular, smooth | Colorless, translucent | Smooth, circular and non-pigmented | Beige, round, slightly convex, regular margins | Colorless, translucent |
| **Cell morphology** | Short to long rods | Short rods | Large rods | Large rods | Short rods |
| **Fluorescent pigments** | nr | + |  | + | + |
| **Motility** | + | + | + | + | + |
| **Growth at 22°C** | + | + | + | + | + |
| **Growth at 30°C** | + | + | + | + | + |
| **Growth at 37°C** | - | - | - | - | - |
| **NaCl tolerance** |  |  |  |  |  |
| **1%** | + | nr | + | + | nr |
| **4%** | + | nr | + | + | nr |
| **8%** | - | nr | - | - | nr |
| **Oxidase** | + | w | + | + | w |
| **Catalase** | + | + | + | + | + |
| **Urease** | - | nr | - | nr | nr |
| **Utilization of:** |  |  |  |  |  |
| **p-Hydroxy-phenylacetic acid** | - | - | nr | + | - |
| **Tween 40** | w | + | nr | - | + |
| **Glycyl-L-proline** | - | - | nr | - | - |
| **Methyl pyruvate** | + | + | nr | + | + |
| **D-Galactonic acid lactone** | + | + | nr | + | + |
| **α-Hydroxy butyric acid** | - | v | nr | + | v |
| **α-Keto butyric acid** | + | + | + | w | + |
| **D-Fucose** | + | nr | - | + | nr |
| **Sucrose** | + | + | + | + | - |
| **D-Rhamnose** | - | nr | - | + | nr |
| **D-Serine** | + | + | - | + | - |
| **Inosine** | + | + | nr | + | + |
| **D-Sorbitol** | + | - | + | + | v |
| **L-Aspartic Acid** | + | - | nr | - | - |
| **L-Glutamic Acid** | + | - | nr | + | v |
| **L-Serine** | + | + | + | - | + |
| **L-Lactic Acid** | + | - | nr | - | - |

+, positive; -, negative; v, variable; w, weak; nr, not reported.

**Table 3.** Comparative physiological characteristics among *Pseudomonas glycinis* sp. nov. PI111 and related type strains. Data from this study and López et al. (López et al. 2012); Pascual et al. (2015); Kwon et al. (2003).

|  | *P. glycinis* sp.<br>nov. PI111<sup>T</sup> | *P. baetica*<br>CECT<br>7720<sup>T</sup> | *P.*<br>*granadensis*<br>278,770<sup>T</sup> | *P.*<br>*koreensis*<br>21318<sup>T</sup> |
| --- | --- | --- | --- | --- |
| **Colony morphology** | White, mucoid,<br>and circular | White,<br>round, and<br>small | White–<br>yellow,<br>circular,<br>convex and<br>mucoid | White-<br>yellow,<br>mucoid |
| **Cell morphology** | Short rods | Large<br>irregular<br>rods | Short rods | Short rods |
| **Growth at 30°C** | + | + | + | + |
| **Growth at 37°C** | - | - | + | - |
| **NaCl tolerance** | + |  |  |  |
| **1%** | + | + | + | + |
| **4%** | + | + | + | + |
| **8%** | - | - | - | - |
| **Motility** | + | + | +, polar<br>flagella | +, polar<br>flagella |
| **Oxidase** | + | + | + | + |
| **Catalase** | + | + | + | + |
| **Urease** | w | - | nr | - |
| **Utilization of:** |  |  |  |  |
| **Tween 40** | w | + | + | + |
| **N-Acetyl-**<br>**glucosamine** | w | + | + | + |
| **Trehalose** | - | - | + | - |
| **D-Galactonic acid**<br>**lactone** | w | + | + | + |
| **D-Galaturonic acid** | - | - | - | v |
| **D-Glucuronic acid** | + | - | - | v |
| **α-Ketoglutaric acid** | + | + | + | - |
| **Glucuronamide** | + | - | + | + |
| **Inosine** | w | + | + | + |
| **Sodium lactate (1%)** | + | - | - | - |
| **Fusidic acid** | + | - | - | - |
| **D-Fucose** | + | - | nr | nr |

+, positive; -, negative; v, variable; w, weak; nr, not reported.

**Table 4.** Comparative physiological characteristics among *Pseudomonas harudinis* sp. nov. SS112 and related type strains. Data from this study and Menéndez et al. (2015); Saati-Santamaría et al. (2018).

|  | *P. harudinis*<br>sp. nov.<br>SS112<sup>T</sup> | *P. abietaniphila*<br>DSM 17554<sup>T</sup> | *P. bohemica*<br>IA19<sup>T</sup> | *P. graminis*<br>PDSM<br>11363<sup>T</sup> |
| --- | --- | --- | --- | --- |
| **Colony morphology** | Non-pigmented, circular, smooth | Clear, translucent, smooth, circular, convex | Round, bright, clear beige, convex and entire | Yellow, glistening, moderately convex, round, entire |
| **Cell morphology** | Short to long rods | Rods | Rods | Straight rods |
| **Motility** | + | + | +, polar flagellum | +, polar flagellum |
| **Growth at 22°C** | + | nr | + | + |
| **Growth at 30°C** | + | + | + | + |
| **Growth at 37°C** | - | - | + | + |
| **NaCl tolerance** |  |  |  |  |
| **1%** | + | nr | + |  |
| **4%** | + | nr | - |  |
| **8%** | - | nr | - | nr |
| **Oxidase** | - | - | + | - |
| **Catalase** | + | + | + | + |
| **Urease** | - | + | - | + |
| **Utilization of:** |  |  |  |  |
| **Glycerol** | + | + | + | - |
| **Rhamnose** | + | + | - | - |
| **Sorbitol** | - | + | - | + |
| **Sucrose** | - | + | - | + |
| **D-Maltose** | - | + | - | - |
| **Malic acid** | + | + | - | + |
| **Galactose** | + | + | - | w |
| **Mannose** | + | + | + | - |
| **Myo-Inositol** | + | + | - | w |
| **Mannitol** | + | + | - | w |
| **Salicin** | - | - | + | - |
| **Melibiose** | - | - | - | + |
| **D-Trehalose** | - | + | - | - |
| **D-Fucose** | + | + | - | + |
| **L-Fucose** | + | + | - | + |
| **D-Arabitol** | + | + | - | + |

+, positive; -, negative; v, variable; w, weak; nr, not reported.

## DESCRIPTION OF *PSEUDOMONAS ARENAE* SP. NOV

*Pseudomonas arenae* (a.re’ nae. L. gen. n. *arenae* of sand).

Motile, rod shape cells (2-4 µm long × 0.8-1.0 wide). Gram-stain negative. Aerobic and mesophilic. Catalase and oxidase positive, urease negative. In addition to the above characteristics, the following carbon sources are used: D-trehalose, sucrose, D-sorbitol, *myo*-inositol, L-pyroglutamic acid, L-serine, D-galacturonic acid, D-glucuronic acid, methyl pyruvate, Tween 40 (weak), γ-amino-butyric acid, β-hydroxy-D,L-butyric acid and propionic acid. The following carbon sources are not used: N-acetyl-D-glucosamine, L-rhamnose, D-aspartic acid, D-serine, Glycyl-L-proline, L-histidine and D-malic acid.

The type strain VK110^T^ was isolated from *Phragmites* sp. in New Jersey, United States.

## DESCRIPTION OF *PSEUDOMONAS GLYCINIS* SP. NOV

*Pseudomonas glycinis* (gly.ci’nis. N.L. fem. gen. n. glycinis of *Glycine max*, the soybean).

Motile, short rod-shaped cells (1-2 µm long × 0.3-0.5 wide). Gram-stain negative. Aerobic and mesophilic. Catalase and oxidase positive, urease negative. In addition to the above characteristics, the following carbon sources are used: L-pyroglutamic acid, L-serine, Tween 40 (weak), γ-amino-butyric acid (weak) and propionic acid. The following substrates do not serve as carbon sources: D-trehalose, sucrose, L-rhamnose, D-sorbitol, *myo*-inositol, D-aspartic acid, D-serine, L-histidine, D-galacturonic acid, D-glucuronic acid and β-hydroxy-D,L-butyric acid.

The type strain PI111^T^ was isolated from *Glycine max* in Missouri, United States.

## DESCRIPTION OF *PSEUDOMONAS HARUDINIS SP.* NOV

*Pseudomonas harudinis* (ha.ru’di.nis. L. gen. n. *harudinis* of reed).

Motile, short to long rods (2-4 µm long × 0.8-1.0 wide) that stain Gram negative. Aerobic and mesophilic. Catalase positive and oxidase negative, with weak urease activity. In addition to the above characteristics, the following carbon sources are used: L-rhamnose, *myo*-inositol, D-aspartic acid, D-serine (weak), L-histidine, D-galacturonic acid, D-glucuronic acid, methyl pyruvate, D-malic acid, bromo-succinic acid, Tween 40 (weak), γ-amino-butyric acid and β-hydroxy-D,L-butyric acid. The following substrates are not used as carbon sources: D-trehalose, sucrose, N-acetyl-glucosamine, D-sorbitol, D-glucose-phosphate, glycyl-L-proline, L-pyroglutamic acid, L-serine and propionic acid.

The type strain SS112 was isolated from *Phragmites* sp. in New Jersey, United States.

## ACKNOWLEDGEMENTS

We would like to thank Professor Aharon Oren, The Hebrew University of Jerusalem, for help with nomenclature.

